# A novel fluorescence and DNA combination for complex, long-term marking of mosquitoes

**DOI:** 10.1101/2020.08.23.262741

**Authors:** Roy Faiman, Benjamin J. Krajacich, Leland Graber, Adama Dao, Alpha Seydou Yaro, Ousmane Yossi, Zana Lamissa Sanogo, Moussa Diallo, Djibril Samaké, Daman Sylla, Modibo Coulibaly, Salif Kone, Sekou Goita, Mamadou B. Coulibaly, Olga Muratova, Ashley McCormack, Bronner P. Gonçalves, Jennifer Hume, Patrick Duffy, Tovi Lehmann

## Abstract

**Background:** Current mark release recapture methodologies are limited in their ability to address complex problems in vector biology, such as studying multiple groups overlapping in space and time. Additionally, limited mark retention, reduced post-marking survival, and the large effort in marking, collection and recapture, all complicate effective insect tracking.

**Method:** We have developed and evaluated a marking method using a fluorescent dye (Smartwater^®^) combined with synthetic DNA tags to informatively and efficiently mark adult mosquitoes using an airbrush pump and nebulizer. Using handheld UV flashlights, the fluorescent marking enabled quick and simple initial detection of recaptures in a simple, field-ready, and non-destructive approach that when combined with a simple, extraction-free PCR on individual mosquito legs provides potentially unlimited marking information.

**Results:** This marking, first tested in the laboratory with *Anopheles gambiae* s.l. mosquitoes, did not affect survival (median ages 24-28 days, *p-adj* > 0.25), oviposition (median eggs/female of 28.8, 32.5, 33.3 for water, green, red dyes, respectively, *p-adj* > 0.44), or *Plasmodium* competence (mean oocysts 5.56 to 10.6, *p-adj* > 0.95). DNA and fluorescence had 100% retention up to 3 weeks (longest time point tested) with high intensity, indicating marks would persist longer.

**Conclusions:** We describe a novel, simple, no/low-impact, and long-lasting marking method that allows separation of multiple insect subpopulations by combining unlimited length and sequence variation in the synthetic DNA tags. This method can be readily deployed in the field for marking multiple groups of mosquitoes or other insects.

## Introduction

Mosquito transmitted malaria kills almost half a million people annually, and mosquito-borne arboviral diseases such as dengue, Yellow Fever, chikungunya, West-Nile virus, and Zika comprise together a further 100,000-300,000 deaths estimated by the WHO annually (www.who.int/news-room/fact-sheets/). While much work has been done over the past century to answer basic questions about mosquitoes such as their population sizes (TAYLOR, TOURE, COLUZZI, & PETRARCA, 1993; Lehmann, Hawley, Grebert, & Collins, 1998; Touré et al., 1998; Athrey et al., 2012), longevity (Polovodova, 1941, 1949; Cook et al., 2006; Krajacich et al., 2017), dispersal patterns and distances (Gillies, 1961; Gillies & Wilkes, 1965; Huestis et al., 2019), and vectorial capacity (Afrane, Little, Lawson, Githeko, & Yan, 2008; Kramer & Ciota, 2015), methodological limitations in tracking wild mosquitoes have left many of these aspects poorly understood. One of the gold-standard approaches for answering such questions is Mark Release Recapture (MRR), which was first applied to mosquitoes over a century ago and has been modified and refined since (Zetek, 1913; Gillies, 1961; Reisen & Aslamkhan, 1979; Service, 1993; Costantini & Della, 1996; Touré et al., 1998; Lehmann et al., 2010; Guerra et al., 2014). All MRR techniques share the same idea: mark multiple individuals, release them, and determine their new location and frequency upon their recapture.

Due to their small size (mostly <5 mm long body and weighing less than 2 mg), marking mosquitoes is limited to light-weight agents. Different types of marking materials have been used in the past including paints/stains (Zetek, 1913; Gillies, 1961; TSUDA & KAMEZAKI, 2013), protein markers (Hagler & Jackson, 2001), powders and dusts (Verhulst, Loonen, & Takken, 2013; Epopa et al., 2017), internal dyes (Bailey, Eliason, & Iltis, 1962), food coloring (Williams, 1962), radio-isotopes (Patterson, Smittle, & DeNeve, 1969; Zhou et al., 2004), and more recently stable-isotopes (Hamer et al., 2012, 2014; Faiman et al., 2019). With advantages to each of the above, many of these marking methods may alter the behavior of the insect (Naranjo, 1990; Dickens & Brant, 2014), shorten its lifespan (Verhulst et al., 2013), wear off over a short period of time (Walker & Wineriter, 1981; Hagler & Jackson, 2001), and have limited diversity of unique tags so only one or few groups of marked organisms can be distinguished (Service, 1993). Through these deficits and the primary implementation of methodologies on local scales, some aspects of mosquito biology such as long-range movement (Huestis et al., 2019) and survival (Lehmann et al., 2010; Dao et al., 2014) have likely been missed, which implies our overall knowledge of basic mosquito behavior is still limited. Additionally, due to the large population sizes of most insects and the labor-intensive nature of the MRR experiment, individuals that are marked tend to be heavily diluted within the un-marked population, limiting recapture.

To address some of the issues described above, we have developed and tested a new marking technique that combines fluorescence and multiple DNA tags that together allow rapid separation of marked from unmarked mosquitoes and near-unlimited mark complexity. Our results demonstrate that it can be efficiently and effectively applied onto large numbers of mosquitoes with a mark that remains detectable for at least 3 weeks with no obvious drop in intensity of both visible and molecular tags. Moreover, experiments demonstrate no impact on mosquito survival, blood feeding, reproduction, and capacity to support *Plasmodium* infection. Thus we find this methodology a straightforward and highly versatile technique well suited to lab and field applications at scale.

## Materials and Methods

### Smartwater^®^ fluorescent spray

For fluorescence marking of mosquitoes we utilized SmartWater^®^ (SmartWater CSI LLC. Telford, UK). SmartWater is a proprietary, traceable water-based solution developed as a forensic marker for concealed labelling of valuable items (see: www.smartwater.com/). Within the SmartWater solution used in our experiments were two components: (1) a non-toxic, water-based fluorescent solution which determines the color of the mark based on different mixtures of the base colors (APEX Invisible Blue, Red, Green, or Cartax DP (Yellow-Green)), rendering the marked object visible under UV light (365nm), and (2) Mowilith LDM 7709, a non-toxic, water-based polymer emulsion used to bind the fluorescence to the marked substrate.

### DNA tag design, sizes and verification

A PCR validated set of 14 DNA tags and primer sequences were created for this project. The tags vary in size from 80 to 340 base pairs (Supporting Information File 1) and share two 20 base universal primer regions allowing for the use of one pair of primers to identify the tag based on size in one reaction mixture. Each tag has a unique internal sequence in addition to its unique length, enabling additional identity confirmation via Sanger sequencing (Sanger, Nicklen, & Coulson, 1977). The sequence of each DNA tag was generated with R (R version 3.6.0) to have roughly a 50% GC content, no start codons, and no significant similarities to known sequences in the nr database of BLAST (Altschul, Gish, Miller, Myers, & Lipman, 1990). Tags were amplified using 1 μl of 0.1 μM single stranded DNA ultramer (Integrated DNA Technologies, Coralville, Iowa, USA) or 1 μl of a 1:100 or 1:200 dilution of a 250ng or 500ng synthesis scale double-stranded DNA gBlock in a 50 μl PCR reaction (GoTaq Green Master Mix, Promega, Madison, WI, USA) with 400nM of primers (IDT). Reactions were amplified at 94°C for 5 minutes, then 32 cycles of 94°C for 30 seconds, 59°C for 30 seconds, 72°C for 30 seconds, and a final elongation of 72°C for 5 minutes. All amplifications were performed in heat-sealed random-access plates (4titude Ltd, Surrey, UK) to reduce contamination risk, and individual-use aliquots were used to add 10 μl of PCR product per 2 ml of SmartWater^®^ spray solution (see Application method below). PCR products were visualized on a 3% agarose / ethidium bromide gel to verify the presence of one band per reaction, and no off-target amplification.

### Application method

Mosquitoes destined for marking were placed in pint-size paper cups with a muslin net cover, onto which the spraying apparatus (Fig. 1) was attached to dispense the spray solution. The spray solution was mixed shortly before spraying onto the mosquitoes as follows: 1-8% fluorescence (color-dependent) were mixed well with 0.2-2.5% polymer, 0.5% synthetic DNA per tag, and topped with deionized water to a final volume of 2 ml. Different colors were found to behave slightly differently when sprayed (i.e. variation in fluorescence adherence to mosquito body parts), requiring adjustment of ratios (see Supporting File 2: SmartWater Recipes). The solution was vortexed at 2000 rpm for 30 seconds to ensure even mixing of the components. Vortexing was repeated immediately before transfer of the marking solution into the nebulizer capsule. The nebulizer set (model NEB KIT 500, Drive DeVilbiss Healthcare, Port Washington, NY) consisted of a capsule (spray solution reservoir, 3 ml in volume), ‘T’-connector (plugged with cotton wool on one end; see Fig. 1), flexible air tube and air supply tube, fed by a 1/5 horse-power airbrush compressor (model TC-20-H6-B, Master Airbrush) preset to 1.5 Bar pressure. A 10-cm diameter powder funnel (model 4252, Thermo Fisher Scientific. Waltham, MA) was attached with its narrow end to the flexible air tube, to allow uniform dispensing of the sprayed solution into the paper pint by covering its top with the conical end, preventing spray from exiting the pint (Fig. 1).

**Figure 1.**
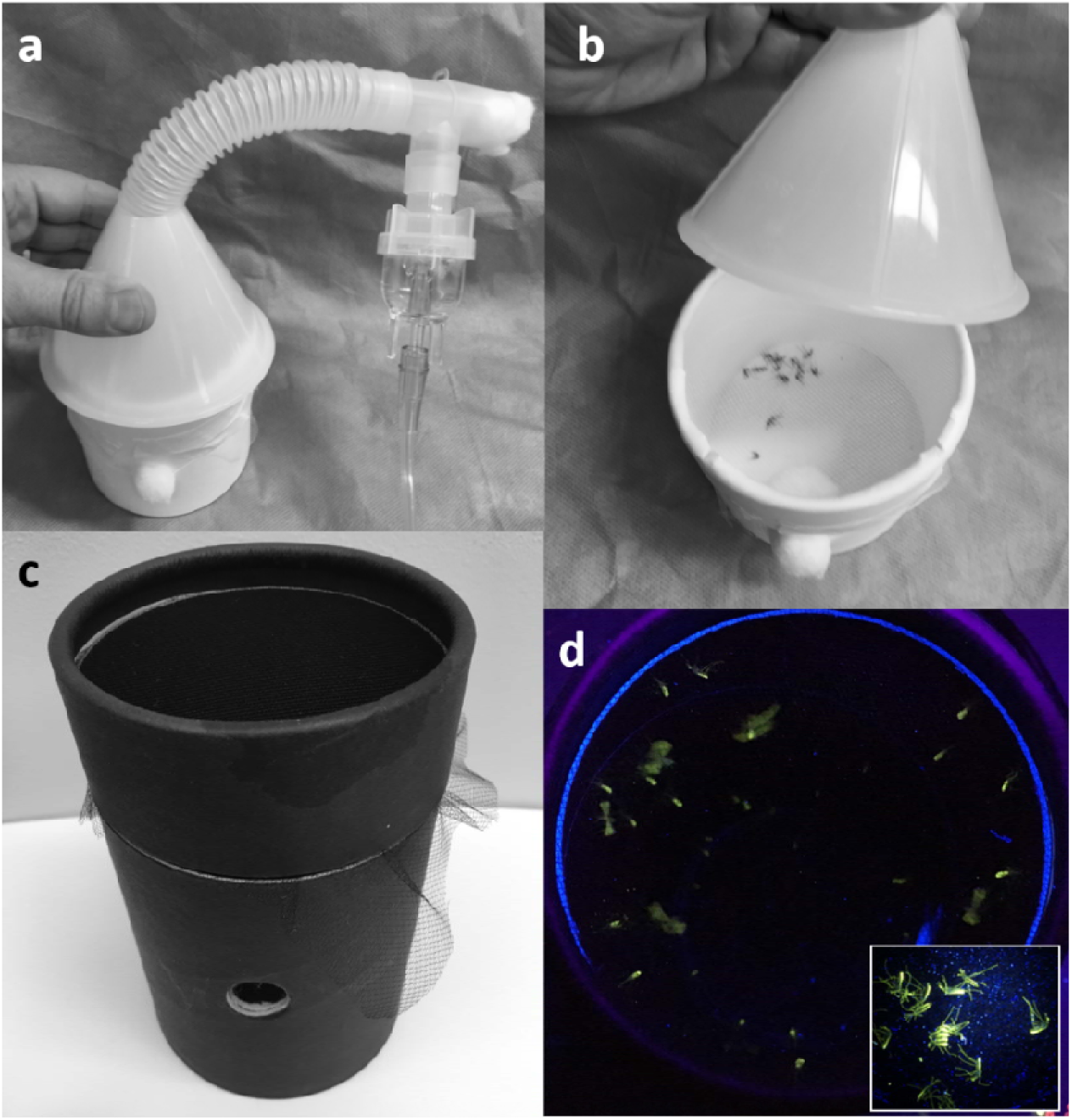
Spray Apparatus: Nebulizer and funnel cone over spray cup (a) with mosquitoes (b). Black paper detection cup (c) which limits background fluorescence and allows for detection of live (d) and dead (d inset) fluorescently labeled mosquitoes using a handheld UV flashlight (365nm).

For spraying, 2 ml of mixed solution was pipetted into the vertically held nebulizer capsule, followed by activation of the compressor for 5-6 seconds, dispensing ~250 μl, with the funnel cone covering the mosquito pint. A visual verification of the spray cloud exudate was done immediately before covering the mosquito pint to rule out the possibility of blockage or non-treatment. The mosquito pint was tapped gently to encourage mosquitoes to take flight during the spraying, ensuring uniform coverage of the mosquitoes by the nebulized spray solution. After 5 seconds of spraying of each pint, mosquitoes were inspected visually. Adequate marking resulted with sedentary mosquitoes, typically seen standing on the cup floor, or flying poorly. Full mobility was regained ~15 minutes after spraying. Sprayed mosquitoes were left in the paper pint no less than 2 hours to allow full drying of the sprayed solution and full recovery of the mosquitoes before subsequent release.

### Effect of SmartWater and DNA spraying on longevity, blood feeding, reproduction, and development of *Plasmodium* parasites

To verify that the application of SmartWater and/or DNA does not affect the survival of mosquitoes, we compared the lifespan of mosquitoes after application of combinations of SmartWater fluorescent components (colors) and polymer concentrations versus a water control. Survival analyses were performed using Kaplan-Meier survival curves and pairwise log rank tests adjusted for multiple comparisons implemented in the ‘survminer’ package in R (Kassambara, Kosinski, & Biecek, 2019). These experiments were performed using *Anopheles gambiae* G3 strain mosquitoes from our insectary at the NIH, or a recently adapted strain *of Anopheles coluzzii* in Mali under standard insectary conditions (27°C, 85% RH, 12:12 Light:Dark cycle). The durability of the SmartWater marking was tested with a simulated “rain” of a weekly water misting of the mosquito rearing cages from a laboratory spray bottle (model F11620-0050, Bel-Art-SP Scienceware^®^, Wayne, NJ). Qualitative intensity of marking post-rain was tracked weekly. A cursory examination of blood feeding and reproduction was also performed, in which groups of 20 mosquitoes were given two bloodmeals and allowed to oviposit, at which point egg counts were estimated per blood fed female. Feeding proportions were compared via an adjusted Dunnett’s test with the ‘binMto’ package in R (Schaarschmidt, 2018).

The potential effects of SmartWater solution on the transmission of *Plasmodium falciparum* parasites were also investigated using a standardized membrane feeding assay (SMFA) following previously published methodology (Wu et al., 2008; Sagara et al., 2018). Briefly, an *in vitro* 15-day old culture of *P. falciparum* (NF54 line) containing stage V gametocytes was diluted with washed O+ RBCs (Interstate Blood Bank, Memphis, USA) and an AB+ serum pool (not heat-inactivated) from US malaria-naive subjects (Interstate Blood Bank, Memphis, USA) to final concentration of 0.07%-0.1% stage V gametocytes and 38.5% hematocrit. For each individual assay, 260μL of diluted culture was fed to groups of ~50 starved 3–5 day old *Anopheles gambiae* (G3 strain) mosquitoes using a Parafilm membrane on a mosquito feeder, kept warm with 40°C circulating water. After feeding, mosquitoes were kept at 26°C and 80% humidity conditions to allow parasites to develop, and survival was tracked daily. On Day 8 after the feed, mosquito midguts were dissected and stained with 0.05–0.1% mercurochrome solution in water for 20-30min. Infectivity was measured by counting oocysts in at least 25 mosquitoes per sample. On Day 15 after the feed, salivary glands of at least 5 mosquitoes per group were dissected and sporozoites were observed by phase-contrast microscopy.

## Results

### Post-mark survival, oviposition, and mark persistence

We tested multiple combinations of fluorescence and polymer components to determine the optimal concentrations in terms of marking visibility under handheld UV flashlight and minimal effect on survival. Different color combinations required varying amounts of polymer for life-long retention on sprayed mosquitoes (Fig. 2). The intensity of the fluorescence varied by color, but at least four colors could provide long-term marking (orange color not shown, Fig. 2). The intensity of marking found with the red mixture was variable and this mixture was not used in later experiments.

**Figure 2:**
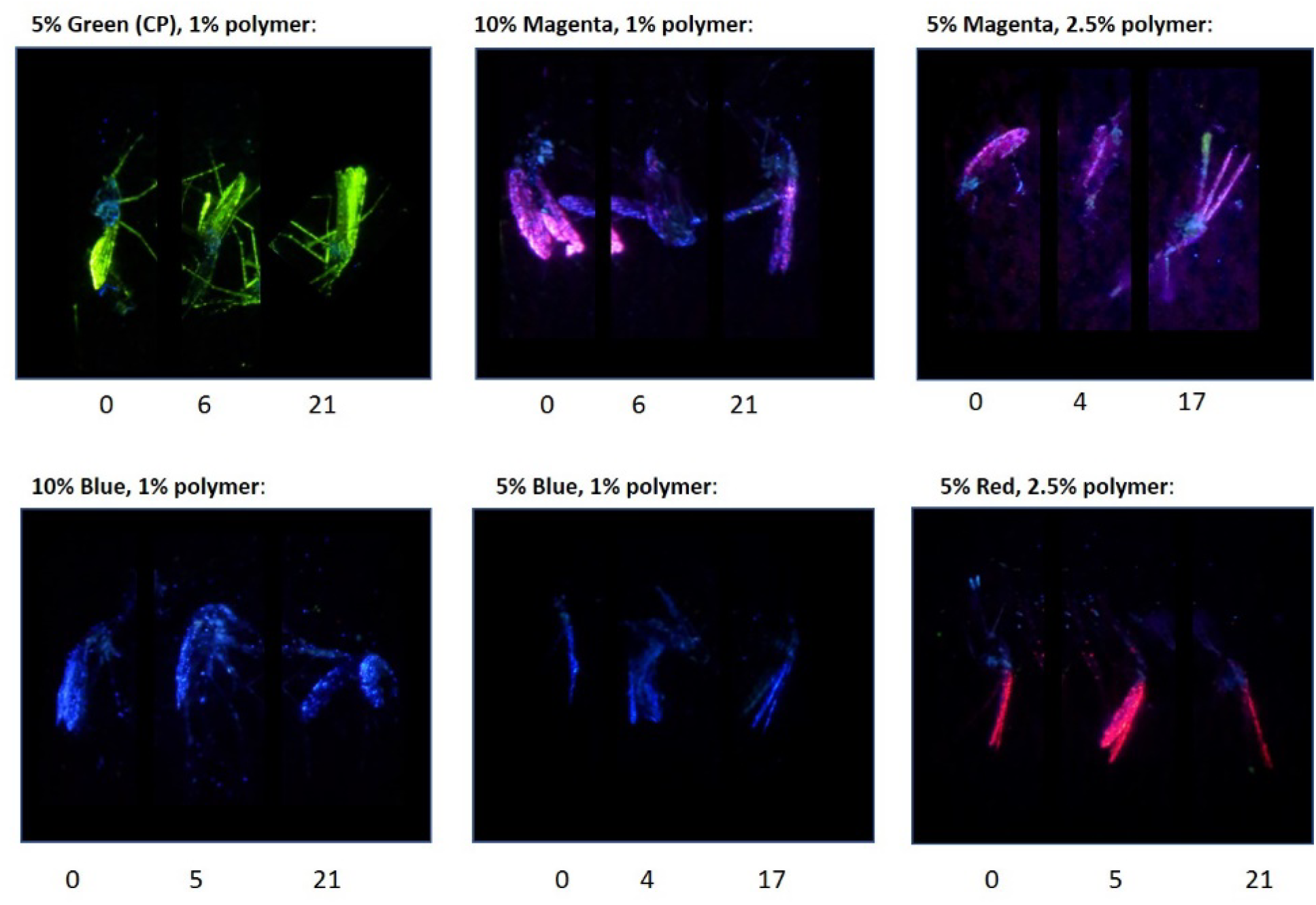
Composite of SmartWater marking intensity (pictures) over time (days post application below each panel).

To evaluate the negative impact that polymer and colors may have on mosquitoes, survival was measured in experiments using Cartax (yellow-green), blue, magenta and orange in comparison with water control (2 replicates ~15 mosquitoes/group, adjusted *p*-values ≥ 0.21, 0.11, 0.96, 0.98 by pairwise log rank test for blue, Cartax, magenta, and orange, respectively). None of the fluorescence and polymer concentrations tested had significant effects on survival (adjusted p-value > 0.48 by pairwise log rank test, Supporting File 3: Mosquito survival per fluorescence/polymer concentration). An early experiment indicated there may be a slight, non-significant increase in mortality at polymer concentrations greater than 2.5%, but this high polymer concentration is not necessary for strong and long term marking.

**Supporting File 3:**
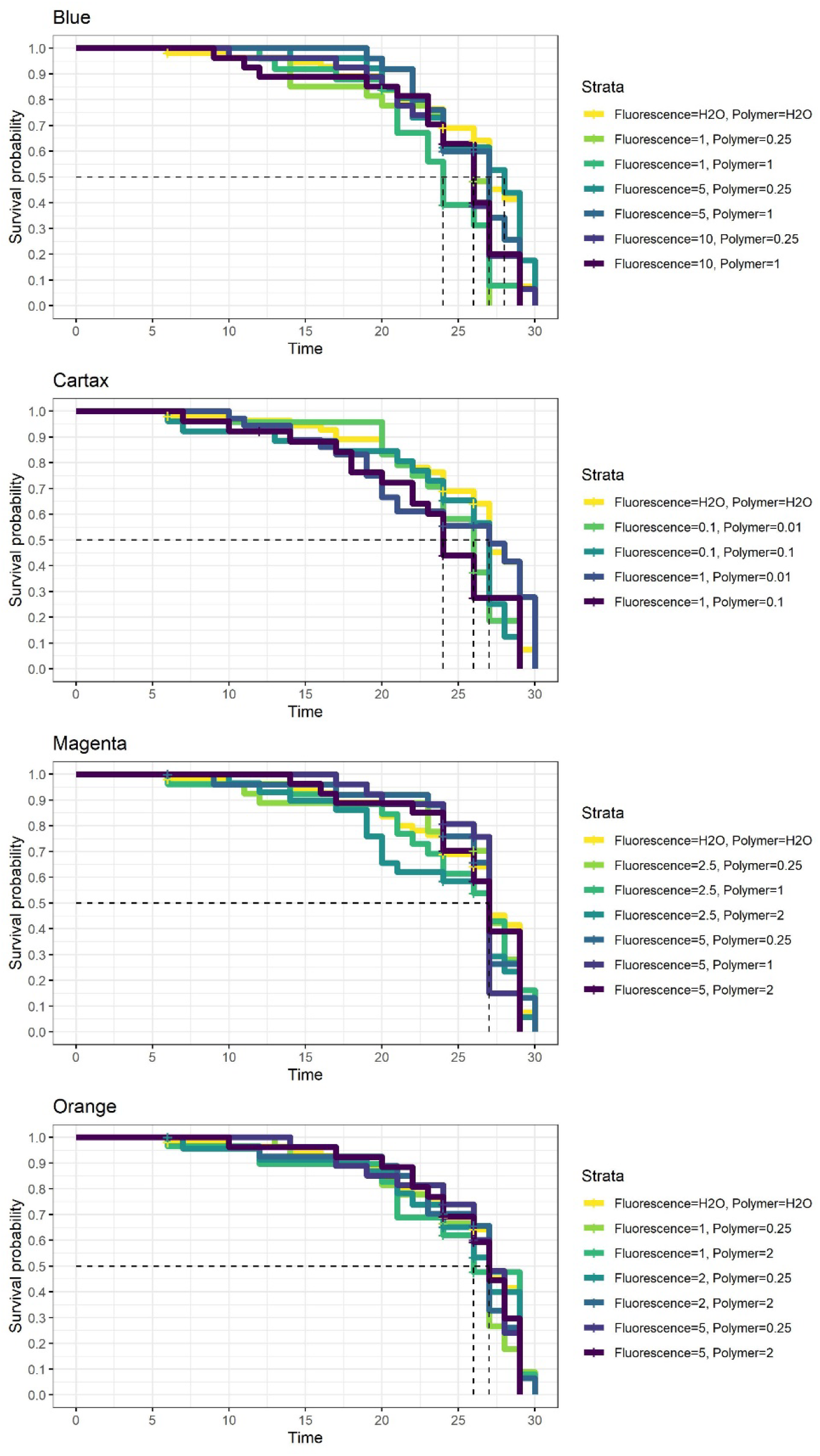
Mosquito survival per fluorescence/polymer concentration: Analysis of the effect of color concentration (fluorescence), and polymer concentration on mosquito survival (days) compared to water control (H_2_O). All comparisons were non-significant (2 replicates, ~15 mosquitoes/group; adjusted *p*-values ≥ 0.21, 0.11, 0.96, 0.98 by pairwise log rank test for blue, cartax, magenta, and orange, respectively). Dotted lines signify when each group reached 50% survival.

In a larger survival study based on the results of the first, no increased mortality was also seen with SmartWater and DNA spray when applied to recently-colonized, laboratory reared *An. coluzzii* mosquitoes in a village kept under insectary conditions in Mali (26-28 °C, 70-85% RH, 2-19 cups per group, 87-1026 mosquitoes, Fig. 3).

**Figure 3:**
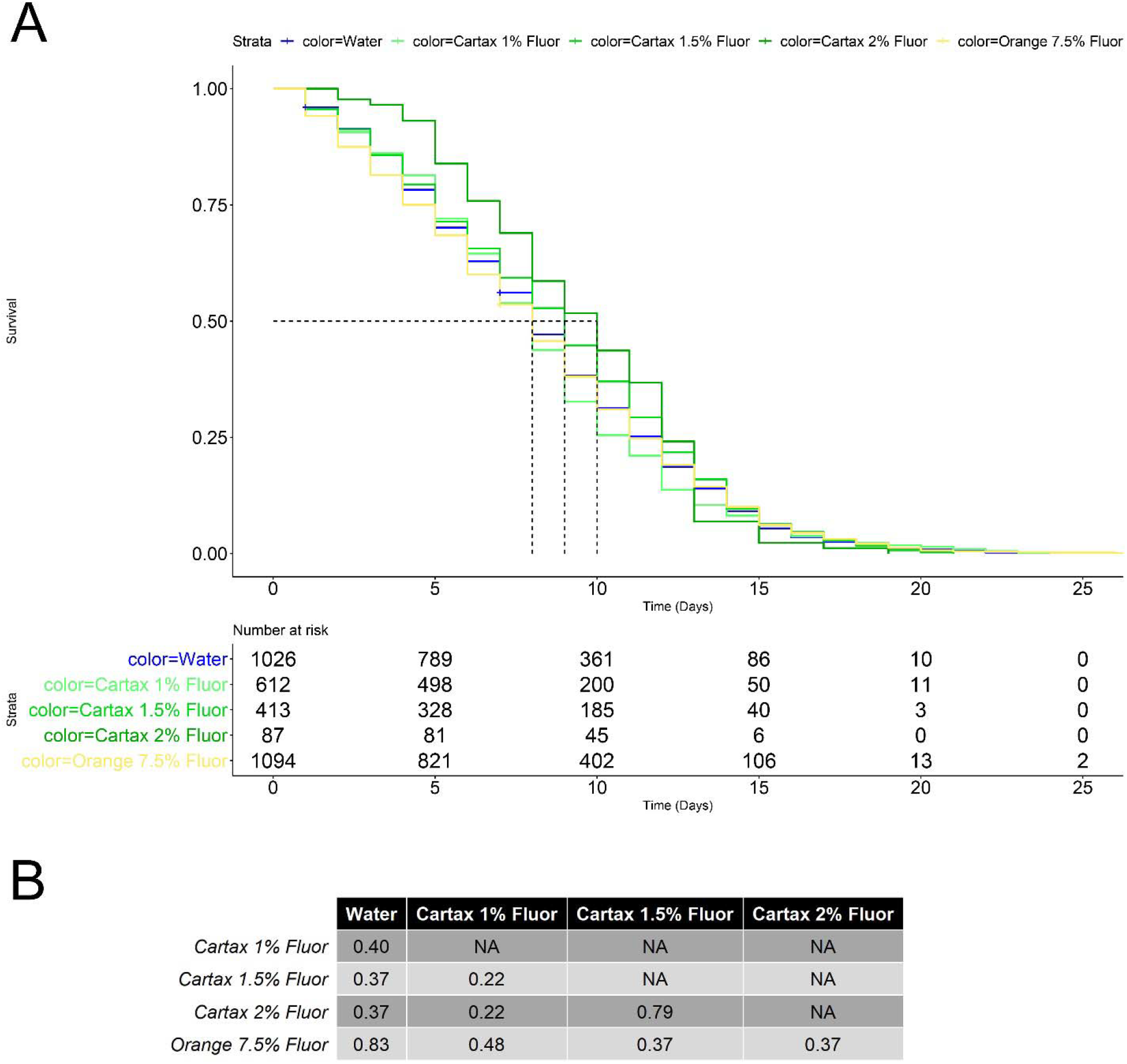
Survival analysis for cartax (green) and orange colors with added DNA tag: Kaplan-Meier survival curves (A-top) and risk table (A-bottom) for 5 variations of colors tested on laboratory mosquitoes in Mali. Non-significant adjusted p-values from pairwise log rank test of survival curves are shown for each comparison (B). Polymer concentrations were 0.3% for Cartax and 2.5% for all other colors.

Additionally, we assessed the application of a weekly simulated rain on one color (Green/’Cartax”). We found that the ‘simulated rain’ treatment had no effect on the visibility of fluorescence on all mosquitoes tested for the duration of the experiment (21 d, data not shown). Finally, feeding and oviposition rates of mosquitoes marked with the fluorescence and DNA mixture did not differ from that of the water-sprayed control using the Dunnett’s test (Supporting File 4: Feeding Rate and Supporting File 5: Feeding and Oviposition, *n*=20/group, 35-90% feeding rate per fluorescence group, 60% water-only control).

**Supporting File 4:**
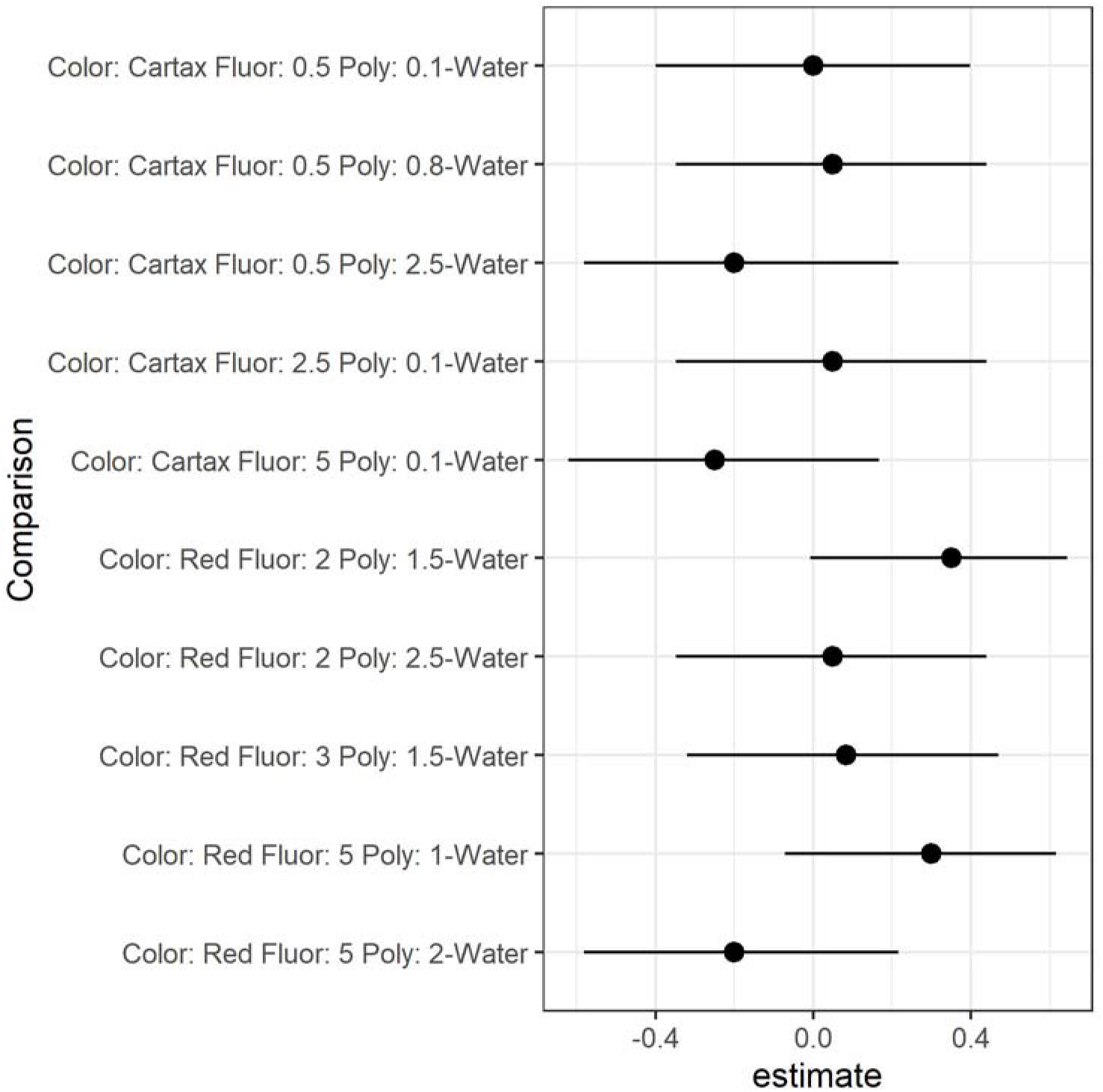
Feeding Rate: Dunnett’s test showing 5-95% confidence intervals of feeding rate of fluorescently labeled *Anopheles gambiae* s.l. compared to control (water). All confidence intervals cross 0 indicating no significant difference in feeding rate to control.

**Supporting File 5:** Table of Feeding and Oviposition: Summary table of feeding and oviposition of *Anopheles gambiae* s.l. mosquitoes sprayed with control (water only) or fluorescence/polymer mixture.

| Color | Fluorescence (%) | Polymer (%) | Females Fed | Total Females | Total Eggs | Average Eggs/Fed Female |
| --- | --- | --- | --- | --- | --- | --- |
| Water | 0 | 0 | 12 | 20 | 345 | 28.75 |
| Cartax | 0.5 | 0.1 | 12 | 20 | 390 | 32.50 |
| Cartax | 0.5 | 0.8 | 13 | 20 | 430 | 33.08 |
| Cartax | 0.5 | 2.5 | 8 | 20 | 30 | 3.75 |
| Cartax | 2.5 | 0.1 | 13 | 20 | 540 | 41.54 |
| Cartax | 5 | 0.1 | 7 | 20 | 25 | 3.57 |
| Red | 2 | 1.5 | 19 | 20 | 500 | 26.32 |
| Red | 2 | 2.5 | 13 | 20 | 575 | 44.23 |
| Red | 3 | 1.5 | 13 | 19 | 517 | 39.77 |
| Red | 5 | 1 | 18 | 20 | 600 | 33.33 |
| Red | 5 | 2 | 8 | 20 | 102 | 12.75 |

### Development of *Plasmodium* parasites in sprayed mosquitoes

The application of SmartWater or DNA had no significant effect compared to no spray controls on the development of *P. falciparum* oocysts at 8 days post feed based on a pairwise t test adjusted for multiple comparisons (mean oocysts 5.56 to 10.6, adjusted *p*-values > 0.95 for all groups, Supporting File 6: Oocyst per mosquito per treatment).

**Supporting File 6:**
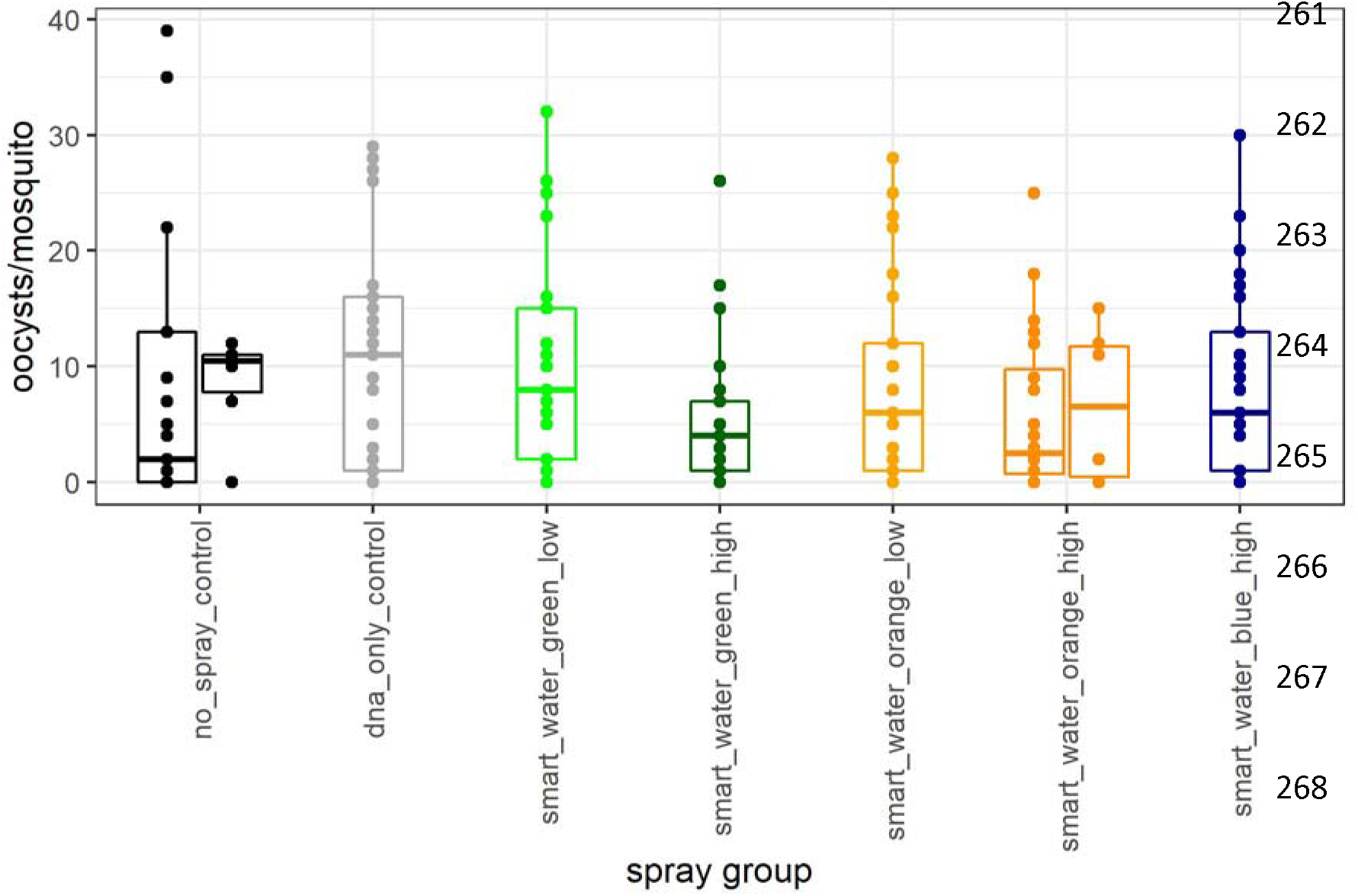
Oocyst per mosquito per treatment: *Plasmodium falciparum* oocysts per mosquito 8 days post standardized membrane feeding assay (n=25 per group). All groups not statistically different by adjusted pairwise dunn test (p-adj > 0.46). Groups are: no spray control (mock application), DNA only control (water with DNA tag), green (cartax) with low fluorescence/polymer (1 and 0.2%), green (cartax) with high fluorescence/polymer (3 and 0.6%), orange with low (7.5 and 2.5%) and high (15 and 3%), and blue with high (20 and 2%) (see Table in Supporting file 7). Multiple replicates per condition are split into individual boxplots.

**Supporting file 7:**
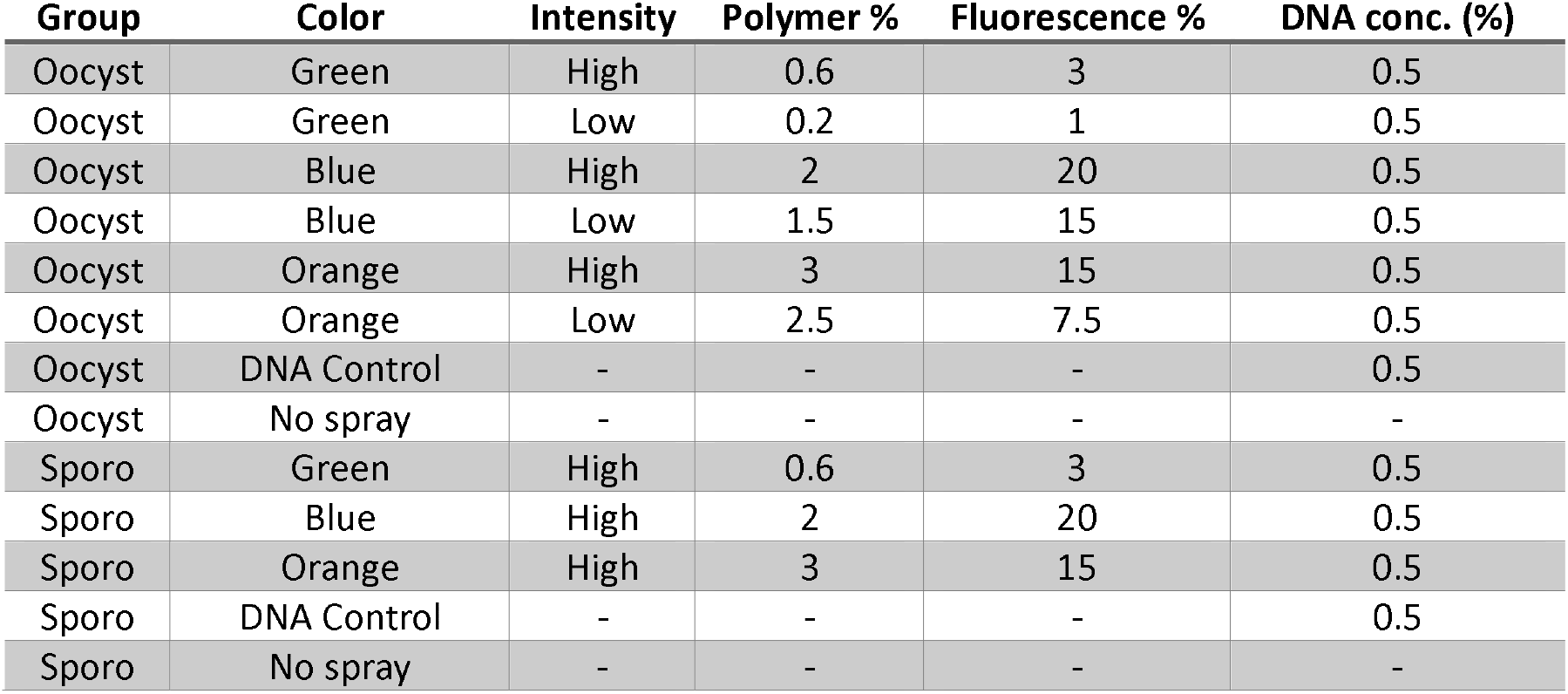
Table of polymer/spray conditions for *Plasmodium* competence experiment.

| Group | Color | Intensity | Polymer % | Fluorescence % | DNA conc. (%) |
| --- | --- | --- | --- | --- | --- |
| Oocyst | Green | High | 0.6 | 3 | 0.5 |
| Oocyst | Green | Low | 0.2 | 1 | 0.5 |
| Oocyst | Blue | High | 2 | 20 | 0.5 |
| Oocyst | Blue | Low | 1.5 | 15 | 0.5 |
| Oocyst | Orange | High | 3 | 15 | 0.5 |
| Oocyst | Orange | Low | 2.5 | 7.5 | 0.5 |
| Oocyst | DNA Control | - | - | - | 0.5 |
| Oocyst | No spray | - | - | - | - |
| Sporo | Green | High | 0.6 | 3 | 0.5 |
| Sporo | Blue | High | 2 | 20 | 0.5 |
| Sporo | Orange | High | 3 | 15 | 0.5 |
| Sporo | DNA Control | - | - | - | 0.5 |
| Sporo | No spray | - | - | - | - |

Mosquitoes for each spray group were also held for 15 days post-feed for detection of sporozoites in the salivary glands. All groups had at least one mosquito (of 5-6 tested) with *P. falciparum* sporozoites in dissected salivary glands, indicating that the application of SmartWater or DNA tags does not prevent the development and potential transmission of malaria-causing parasites.

### Durability of synthetic DNA as a high-information marker

We tested the incorporation of synthetic DNA tracer tags into the nebulized SmartWater mixture to provide increased group resolution e.g., different tags for mosquitoes released from different release zones and at different dates into the marking mixture. We designed a panel of 14 size-discriminative DNA tags which can be amplified via one pair of primers (Supporting File 8: DNA tag sizes).

**Supporting File 8:**
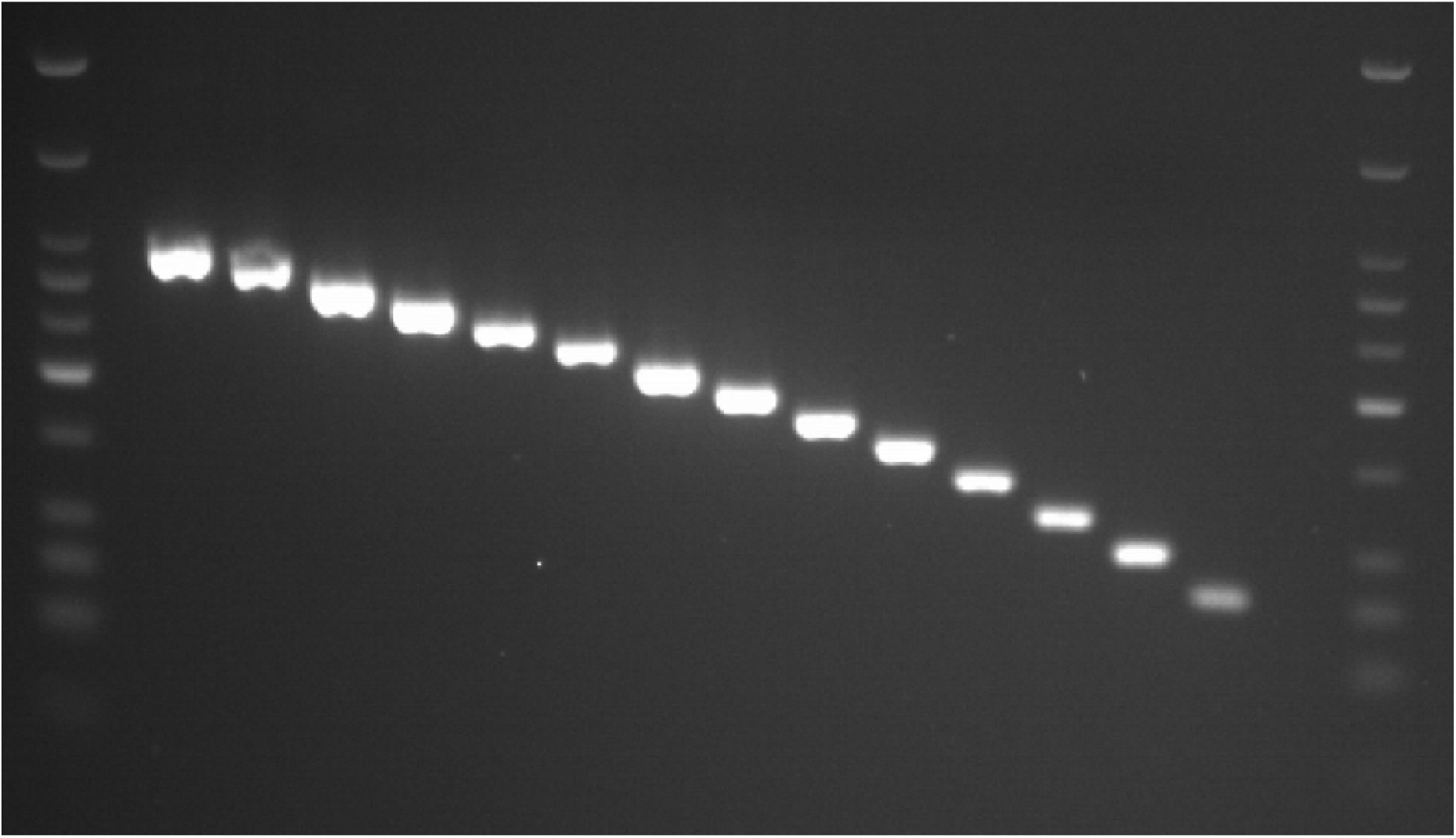
DNA tag sizes: Size based discrimination of DNA tags. Lanes 2-15 DNA amplicons from 340 to 80 bp in 20 bp increments. Lanes 1 and 17: NEB Quick-load low-molecular weight ladder.

For initial testing of DNA tags, we chose one size (120 bp) to be incorporated with SmartWater solution and track through the mosquito lifespan for detection, though detection of multiple sized amplicons on mosquitoes is possible (up to four tags with two primer sets on one mosquito has been tested, data not shown). To reduce costs and time needed for DNA extraction, we evaluated the ability of direct PCR detection by using a portion of the mosquito and adding it directly into the PCR mixture for amplification. We found that mosquito legs provided a reliable PCR-based detection of the sprayed tag over the course of the mosquito’s lifespan without obvious drop in band intensity up to 21 days post spray application (longest period tested, Supporting File 9: Body Part DNA Retention).

**Supporting File 9:**
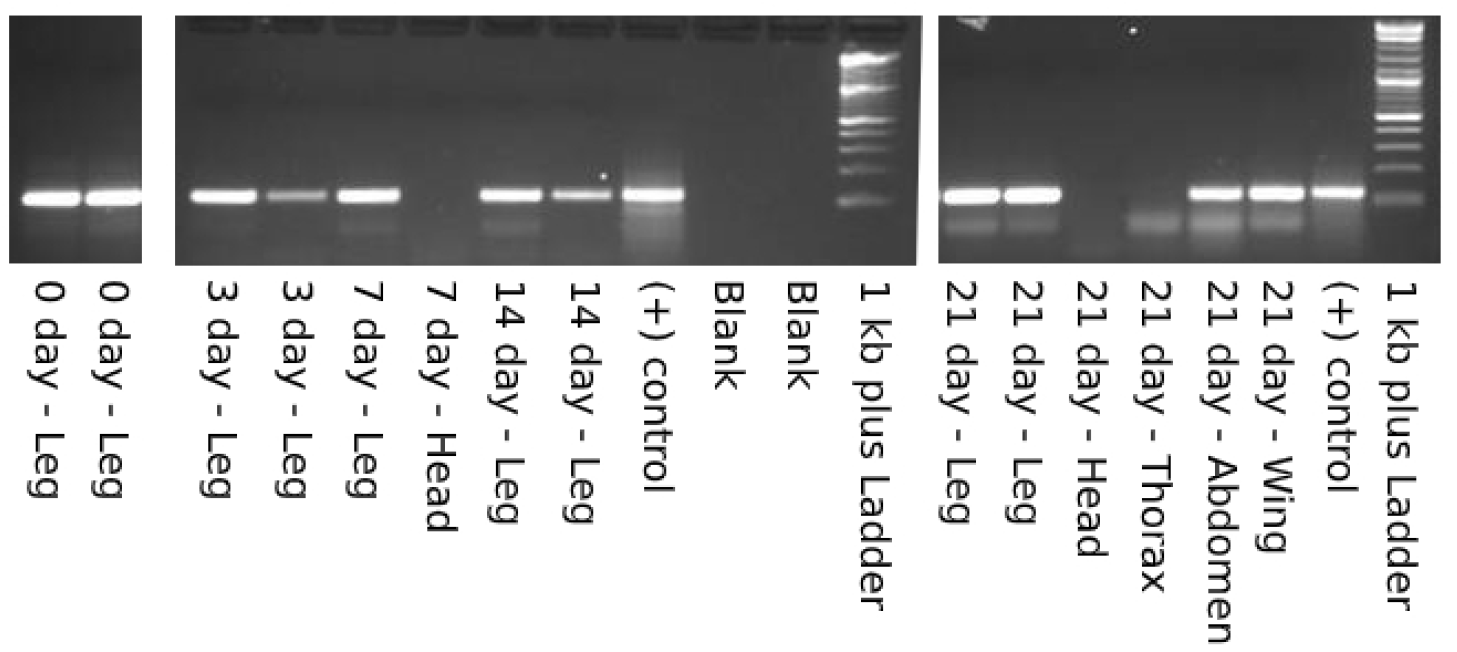
Body Part DNA Retention: Retention of DNA tag over time by body part. Top band of 120 bp shows DNA tag utilized in experiment, sub-band of ~40bp is a primer-dimer. Lanes 1 and 2 were run with the samples displayed in the last panel of the figure. Bottom bands of the ladder used are 100, 200, 300, 400, and 500 from bottom up, respectively.

Similarly, in a second experiment performed on colony mosquitoes in Mali no obvious drop in band intensity was found up to 19 days post spray application (Fig. 4). Wings and abdomens also provided life-long DNA-based detection (Fig. 4).

**Figure 4:**
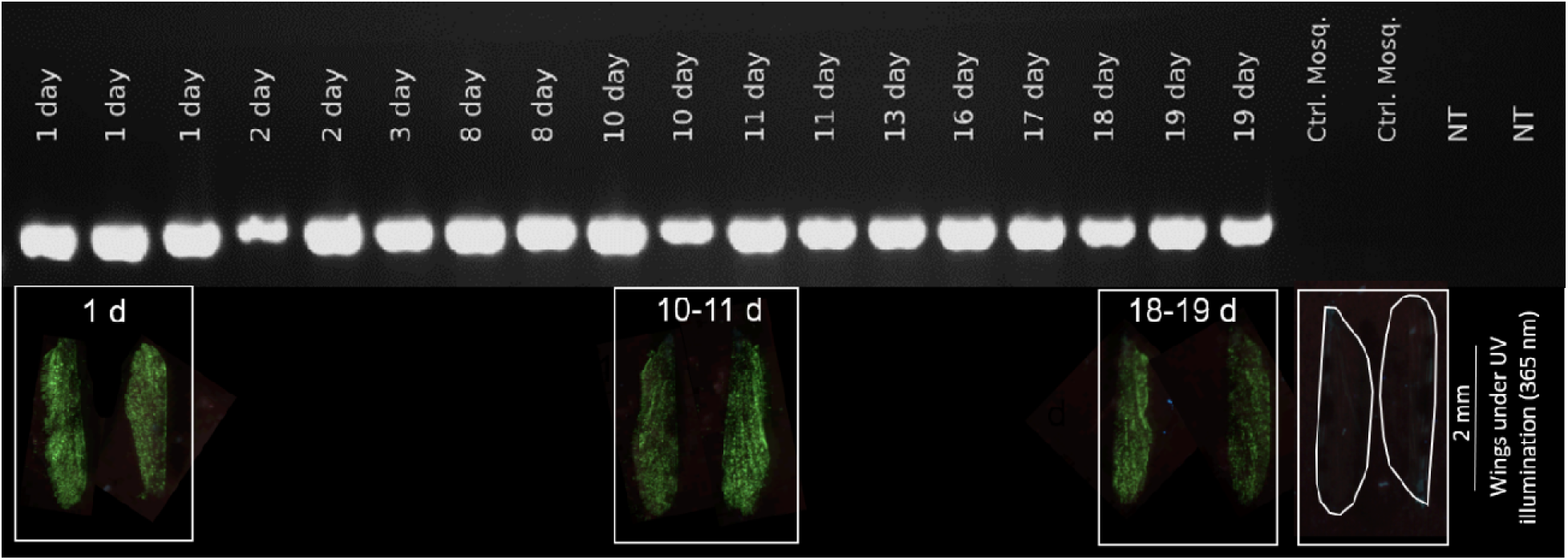
Leg PCR (DNA) and Wing Fluorescence Retention: 220 base pair DNA tag directly amplified from the leg of laboratory reared *Anopheles coluzzii* upon death (lanes 1-18, day of death post marking listed). Two unsprayed control *Anopheles gambiae* (lanes 19-20) and two no template (NT) PCR controls (lanes 21-22) are included as negative controls. Wings of corresponding *A. coluzzii* mosquitoes under UV illumination (365 nm) placed below DNA bands to illustrate visual detectability by duration post treatment in days (all wings from same spray treatment group; two wings selected in random per age group). No significant reduction in intensity of UV was found among the different ages (see Supporting File 10).

**Supporting File 10:**
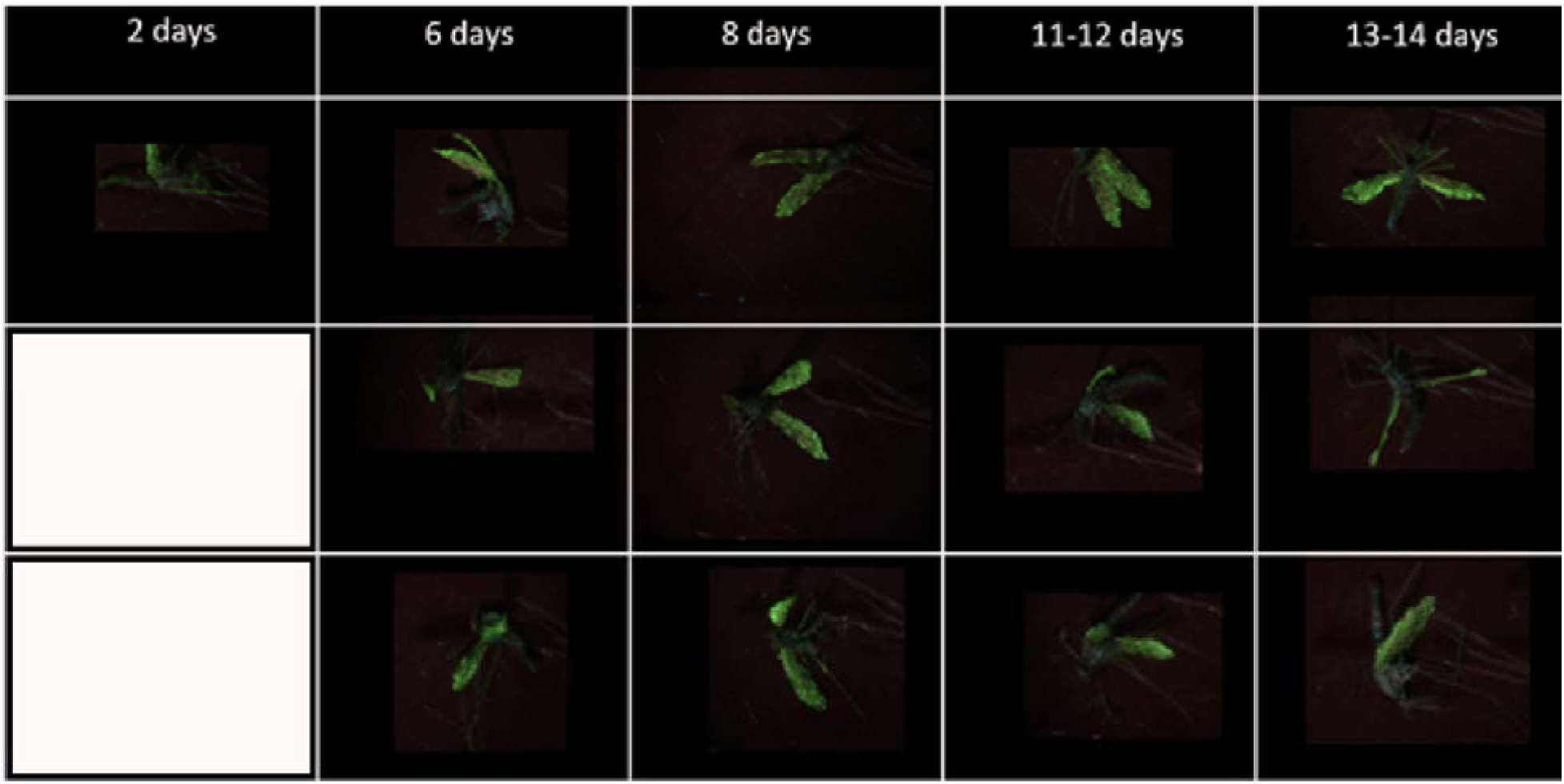
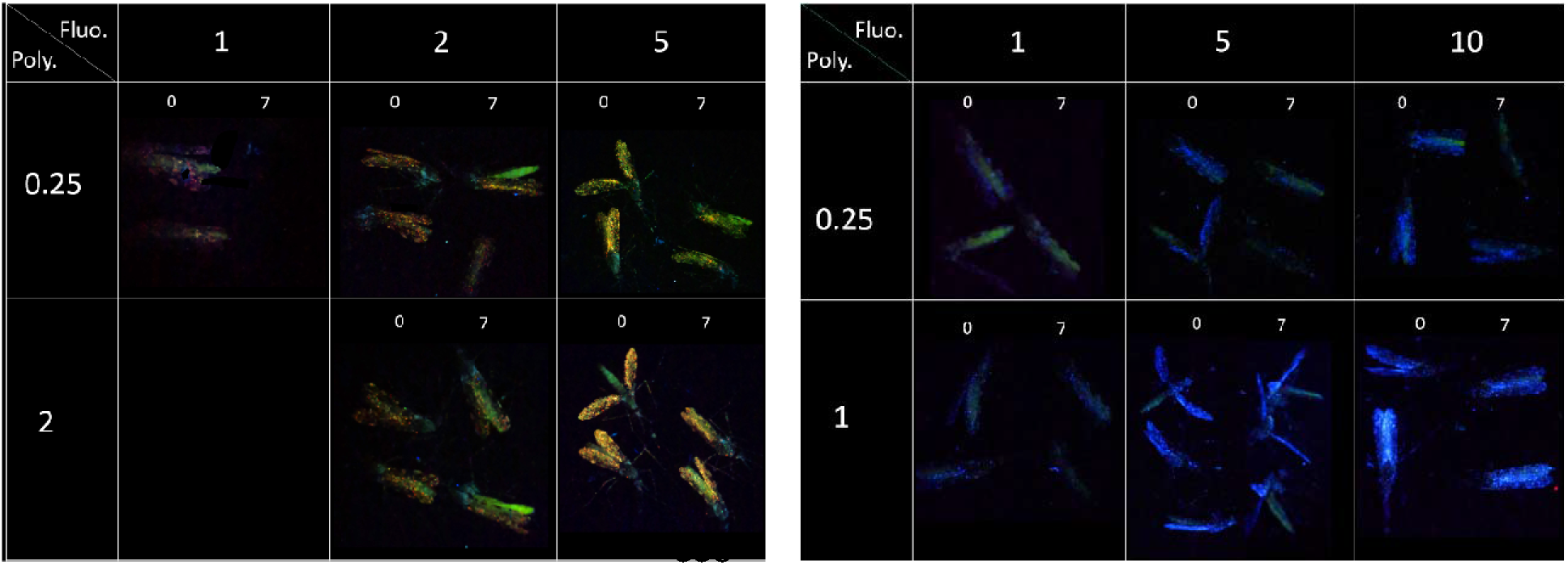
Whole body mosquito fluorescence intensity at death for cartax (top), orange (bottom left), and blue (bottom right) sprayed mosquitoes. Only one mosquito was imaged for Day 2 of cartax-spray mosquitoes. Wings of green spray group from Day 1, 10-11 and 18-19 appear in DNA gel (see Fig. 4).Bottom panels contain multiple polymer (rows [%]) and fluorescence (columns [%]) ratios imaged on day of spraying and seven days after spraying. Note yellowish hue under 1% fluorescence is mosquito abdomen auto-fluorescence.

## Discussion

While the widespread utilization of fluorescent powders and dyes for mark-recapture studies of insects has produced valuable body of information over the past decades (Guerra et al., 2014), methodological limitations leave considerable room for improvements. We investigated the use of synthetic DNA tags co-sprayed with SmartWater^®^ for mosquito marking. Our results demonstrate that the fluorescent mark allows rapid identification of recaptures and the multitude of possible DNA tags facilitate complex experiments by marking with unique tags multiple groups, for example several release sites, various dates, indoors vs. outdoors caught wild mosquitoes, laboratory raised vs. wild mosquitoes, and their combinations. Moreover, we show that DNA amplified directly from legs without extraction, produced bright, life-long detection (up to 3 weeks, highest value tested) with no loss of signal (Fig. 4). Synthetic DNA has been used as an environmentally non-harmful tracer of groundwater sources for several decades in hydrological studies (Sabir, Haldorsen, Torgersen, & Aleström, 1999), and more recently similar DNA barcodes have been developed for safe labeling in food production (SafeTraces, Pleasanton, CA). These DNA tags expand experimental possibilities, and shift the decision about the number of tags to logistical considerations, including the ability to amplify and apply many unique tags for multiple groups per day. Combining the four different colors that were easily discriminated under lab and field conditions by our team (Fig. 2) with the 14 validated DNA tags used in this study allows for up to 56 unique marking groups with a relatively simple experimental design and modest cost. The size-based discrimination of tags with shared primers coupled with Sanger sequencing of the unique internal sequences allows for a blend of rapid detection and more stringent confirmation of tag identity needed for certain field studies.

The use of SmartWater^®^ based fluorescent dyes co-applied with polymer were chosen assuming the mark would be less hazardous, more durable and would minimize the risk of transfer between insects during mating or co-housing. SmartWater conforms with the five prerequisites of an ideal marking material as was defined by Hagler and Jackson (Hagler & Jackson, 2001): highly durable (>30 day retainment in laboratory study), inexpensive (currently supplied gratis by SmartWater Foundation), nontoxic, easily applied, and clearly identifiable. Detection of marked mosquitoes in the field can be achieved through use of a hand-held UV flashlight, easily allowing separation of marked from non-marked individuals following minimal training. With this technique we did see a slight drop in fluorescence over the lifespan of the mosquitoes (Fig. 4, Supporting file 10), however this drop in fluorescence does not appear to be significant enough to limit detection over a likely wet season mosquito lifespan.

DNA tags amplified from ultramers (200bp or less) or gBlocks (>200bp) have limited cost, with one ~$100 tag allowing for preparation of enough spray material for 40,000 spray applications on cups of up to ~50 mosquitoes. Design, amplification, and detection of novel DNA tags requires lab-standard molecular biology knowledge (i.e. primer design, checking BLAST for uniqueness of sequences, etc.), and a thermocycler/gel electrophoresis equipment now standard in many labs, including entomological labs in malaria endemic regions. Spray application only requires an air compressor (~$60-100 depending on model), power source (i.e. 12V adaptor in car), and disposable nebulizer kits (<$3 per nebulizer) which allow for marking thousands of mosquitoes per day depending on number of marks used and team size.

Potential downsides to this application method were found through initial laboratory testing and mock implementation in the field. It was found that significant care had to be taken to use best laboratory practices to prevent cross-contamination as minute amounts of DNA could produce a faint band. As the approach is essentially a nested PCR that is known for high sensitivity and possibility of false positives, this result is not unexpected but must be accounted for. This means that all spray aliquots of the DNA tags should be prepared in a designated area and amplified in heat-sealed individual use tubes to prevent any possible cross-contamination after amplification. At a minimum, the use of pre- and post-amplification areas, the use of one-directional workflow (i.e. do not go from post-amplification to pre-amplification areas), single use reagent aliquots, and use of designated pipettes and lab coats are recommended to limit these issues (Aslanzadeh, 2004). Furthermore, mosquito leg testing should always be done with proper negative control legs to indicate whether there is potential cross-contamination. In the studies in Mali, we found that bright external fluorescence and bright electrophoresis bands present on a mosquito was a reliable indication of true marked mosquitoes. However, it was common for there to be small particles of fluorescence from potentially environmental particles (Reeves, Brookman, & Hammon, 1948), autofluorescence of sugar meals, eggs and/or genitalia (see Supporting File 10), and low degree of contamination from re-used spray cups/covers and aspirators used to process many marked mosquitoes. Care must be taken to not re-use items exposed to many marked mosquitoes, and it is important to separate between spray areas (stations) using different DNA tags and separate them from detection stations. We suggest the use of a visual reference for strong marking such that mosquitoes viewed under microscopy should have >1000 spots of fluorescence per wing, >40 on the head/thorax, and >50 on the legs. Mosquitoes with less than 50 total spots of fluorescence should be treated as potential false positives, though these are rare when taking precautions. Finally, the method presented here still demands time-consuming manual application of the mark on the collected/reared mosquitoes by spraying groups of mosquitoes. Future works would benefit from self-marking techniques which could potentially increase the output of marked individuals several-fold with little-to-no increase in labor. Increasing the numbers of marked individuals significantly may open the door to studies on greater spatial scales as well.

## Conclusions

The SmartWater and DNA tagging mixture provided a highly flexible, modestly priced, and long-lasting mark that had no detectable effects on mosquito survival, despite exceptionally large experiments to evaluate such effect. Additionally, no effect was detected on blood feeding, reproduction, or malaria-transmission capacity. This technique is well-suited for use in studies on mosquito behavior, population size estimation, movement analysis, transmission dynamics, and impact of control measures, especially when multiple groups require higher spatial resolution than is available with currently used adult-marking methodologies. Currently, large MRR experiments using this novel marking method are being performed in the field in Mali showing promising results.

## Acknowledgements

SmartWater CSI^®^ and The SmartWater Foundation for providing materials and aid in optimizing formulations. Andre Laughinghouse, Kevin Lee, and Sam Moretz for logistic support. Peter Tung for NIH legal support and MTA. Drs. Cheick Traore and Laure Juompan provided vital assistance in field operations. We thank the residents of the village Doneguebougou and Thierola for their cooperation and hospitality. This study was supported by the Division of Intramural Research, National Institute of Allergy and Infectious Diseases, National Institutes of Health, Bethesda MD.

## Authors’ contributions

RF, BJK and TL conceived the ideas and designed methodology; RF, LG, BJK, AD, ASY, OY, ZLS, MD, DS, OM, AM, DS, MC, SK, SG, MBC and TL collected the data; BJK, RF, BPC, OM, AM and TL analysed the data; RF, BJK and TL led the writing of the manuscript. All authors contributed critically to the drafts and gave final approval for publication.

