## Supplemental information for "A novel fluorescence and DNA combination for complex, long-term marking of mosquitoes"

**SmartWater CSI with DNA tag – Mosquito marking protocol**

**Preparing CARTAX (Green-Yellow) Smartwater DNA tag solutions and spray applications**

**** All pipetting must be using calibrated and recently tested (within 2 months) pipettes by an experienced technician who has worked with viscous solutions. After vigorously inverting the source solution 7 times, the technician should slowly aspirate from the middle of the volume (not the foamy top), confirm the volume by pipetting at least 2-3 times to see that the solution reaches the same volume and the tip is not clogged. Also that there is no air in the pipette tip before transferring the volume to new tube and confirming no residual fluid in the tip before ejecting the tip.***

Ingredient	volume (ul)	Comment	[Final vol spray sol= 2 ml [change as needed]

CARTAX Green – 1.5%Fluores + 0.3%Pol Final Vol=2ml

SmartWater Cartax Fluorsecense Sol.	30 ul

SmartWater Polymer	6 ul

Location DNA marker (L series)	10 ul *see map

Date DNA marker (A or B series) –	10 ul

[Outdoors DNA marker (L series): optional	10 ul * select the correct one for the group needed]

Deionized (or clean tap) Water	1,944 ul	-without the outdoor marker [1958 ul with it]

MIX well each solution by pipetting up/down x5 with water volume. ALSO mix by inversion before use.

ORANGE – 7.5%Fluores + 2.5%Pol Final Vol=2ml 🡨🡪 Orange Fluores. =1.5 red +0.5 green <*not* Cartax>

SmartWater Orange Fluorsecense Sol.	150 ul

SmartWater Polymer	50 ul

Location DNA marker (L series)	10 ul * see map

Date DNA marker (A or B series) –	10 ul

[Outdoors DNA marker (L series): optional	10 ul * select the correct one for the group needed]

Deionized (or clean tap) Water	1,780 ul	-without the outdoor marker [1770 ul with it]

MIX well each solution by pipetting up/down x5 with water volume. ALSO mix by inversion before use.

Magenta – 7.5%Fluores + 2.5%Pol Final Vol=2ml

SmartWater Magenta Fluorsecense Sol.	150 ul

SmartWater Polymer	50 ul

V/P Location DNA marker (L series)	10 ul * see map

Date DNA marker (A or B series) –	10 ul

[Outdoors DNA marker (L series): optional	10 ul * select the correct one for the group needed]

Deionized (or clean tap) Water	1,780 ul	-without the outdoor marker [1770 ul with it]

MIX well each solution by pipetting up/down x5 with water volume. ALSO mix by inversion before use.

**SPRAY PROCESS**

Use cleaned capsule, T connector, funnel, and plastic tubes for each day/solution. If you change a solution with a different DNA (or fluorescence) tag -Change capsule, T connector, tubes and funnel.

Estimate the volume of solution you need to spray based on the number of houses you have mosquitoes from (250 ul /cup; so 10 cups need 2.5 ml and 40 cups 10 ml). During the first 10 operations, we will also tag 80-100 mosquitoes using 4 cups each with 20-25 mosquitoes). So please add the 4 cups.

1. Check the pumps works and its air pressure is on 1.5 bars by blocking tube tip to allow pressure to build. Adjust to 1.5 bars using rotating knob and lock into lock position when set.

2. Set up ~10 cups in a clean tray and arrange in 2 rows. The cups must be after screening to remove recaptures and must be labeled from houses in the Marking Area. Arrange all cups in trays for quick spray application.

Add up to 4-10 ml

spray solutions are pipetted/poured into nebulizer capsule near external wall.

3. Cover capsule with cap and twist to secure position; Attach to pump output tube to the bottom of the nebulizer/capsule, plug larger hole on the T connector with cotton ball.

4. Holding the nebulizer vertically, connect the flex wide-spray-tube to the T connector and to the funnel

5. Place the connected funnel over the first cup, turn on the pump, and observe a few seconds to see the mist. Once you see the mist hold the funnel close to cover the full opening for 4-5 seconds (look on Timer) while tapping the cup to force the mosquito to fly. Their flight is important. Move to the next cup

6. Without stopping start immediately to spray while tapping for another 4-5 seconds, and so on, until the last cup. Turn off the pump, or move to the next tray if there are more cups that were prepared.

7. Observe mosquitoes immediately after each cup is sprayed. They will be mostly on the floor or standing in the remaining mist for <10 min. They should not show any effects after 15 min and mortality should be same as controls (test will be carried out)

8. Nebulizer parts, from funnel to capsule should be immediately cleaned: 1^st^ by removing the spray washed in running water and then connecting them with the capsule filled with clean water and running the pump for 0.5 minute. Then, wash again and place in a dilute bleach solution for 6-24 hrs. Wash again with clean water after the bleach and let dry in clean place >5m away from the spray station.

9. Do not aspirate or disturb the mosquitoes sprayed for >2 hr (ideally 3 hrs).

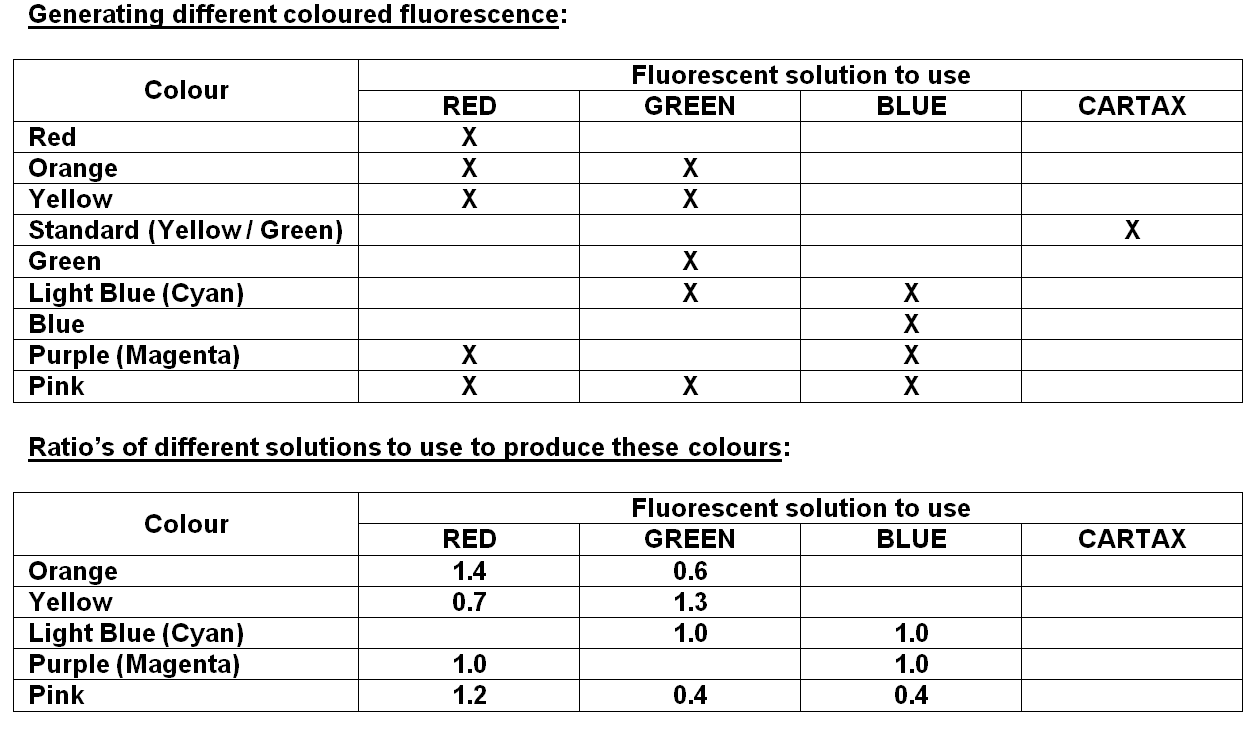
